# First Glimpse of Gut Microbiota of Quarantine Insects in China

**DOI:** 10.1101/2020.12.10.420612

**Authors:** Yanxue Yu, Qi Wang, Ping Zhou, Na Lv, Wei Li, Fangqing Zhao, Shuifang Zhu, Di Liu

**Author notes:** Corresponding authors. (DL), and (SZ). Contributed equally (Yanxue Yu, Qi Wang).

## Abstract

Quarantine insects are economically important pests that frequently invade new habitats. A rapid and accurate monitoring method to trace the geographical source of invaders is therefore needed for prevention, detection, and eradication. Current methods based on insect genetics are often too slow. We developed a novel tracing method based on insect gut microbiota. The source location of microbiota of insects can be used to rapidly determine the insects’ geographic origin. We analyzed 179 gut microbiota samples belonging to 591 individuals of 22 quarantine insect species collected from 36 regions in China and abroad. The gut microbiotas of these insects mainly included Actinobacteria, Bacteroidetes, Cyanobacteria, Firmicutes, Proteobacteria, and Tenericutes. The diversity of the insect gut microbiota was closely related to geographic and environmental factors. Different insect species could be distinguished at the phylum level of microbiota. Populations of individual insect species from different regions could be distinguished at the genus level of microbiota. A method for determining the geographical origin of invasive insect species was tentatively established, but its practical applicability requires further study.

## Introduction

Insects are common, diverse, and widely distributed throughout the world [1]. Quarantine insects are introduced species that may cause harm to agriculture, forestry, stored products, or human health. Countries or regions must take preventive and control measures to reduce the damage caused by quarantine insects. Globally, quarantine insects cause annual losses of billions of dollars [2,3], and China is seriously affected by many species of quarantine insects [4,5]. There are 150 species or genera of quarantine insects on the List of Imported Plant Quarantine Pests in the People’s Republic of China (http://www.aqsiq.gov.cn/xxgk_13386/zvfg/gfxwj/dzwjy/201706/t20170614_490858.htm). Some quarantine insects have a relatively limited distribution. For example, *Leptinotarsa decemlineata* is mainly distributed in the northeast and northwest of China, while other species such as *Lissorhoptrus oryzophilus* are widely distributed in areas where its host plant (*Oryza sativa*) is grown.

Rapid and accurate methods are needed to identify geographical sources of quarantine insects. This would aid in detection, monitoring, prevention, and eradication. Current methods mainly use genetic tests such as DNA barcode technology [6,7], restriction fragment length polymorphisms (RFLPs) [8], single nucleotide polymorphisms (SNPs) [9], and microsatellite markers [10]. All of these methods are based on gene flow between populations, and they require several generations of insects to complete. The genetic changes in populations from different areas can only be used to infer the movement routes [5]. Currently, there are no methods that can trace the origin of an individual insect within a short time interval.

Gut microbiota are often considered to be the second genome of insect species. Gut microbiota are closely linked to the insect’s environment; as such, they are variable and can change rapidly. Microbe characteristics are related to their area of origin; specific microbes can interact with quarantine insects through the diet and thereby contribute to the insect gut microbiota. Gut microbiota might thus provide clues to the geographical source of a quarantine insect. The 16S rDNA gene sequencing technique can be used for detection and identification of insect gut microbiota [11]. In this study, we developed a method to determine the geographical source of quarantine insects using gut microbiota identified by 16S rDNA gene sequencing.

## Results

### Sample collection and DNA extraction

A total of 591 quarantine insect individuals were collected. There were 22 species belonging to 19 genera and 13 families (Table S1). All insects were analyzed using mitochondrial cytochromec oxidase subunit I (COI) genes for species classification (**Figure 1A**). Photos and sizes of the species are shown in **Figure 1B**. The 36 collection sites were located in eight provinces and covered northeast, central, south, and northwest China (**Figure 1**, Table S1). Among the quarantine insects studied, *Ips typographus* was widely distributed in China and has been found in at least 20 provinces. The distributions of the other 21 species varied, and each was only found in several provinces. In addition to the insects collected in China, we also sampled 30 *Ips typographus* and five *Cydia pomonella* individuals that originated outside of China and that were alive when they were intercepted at the ports. In general, these insects are widely distributed and established in China.

**Figure 1.**
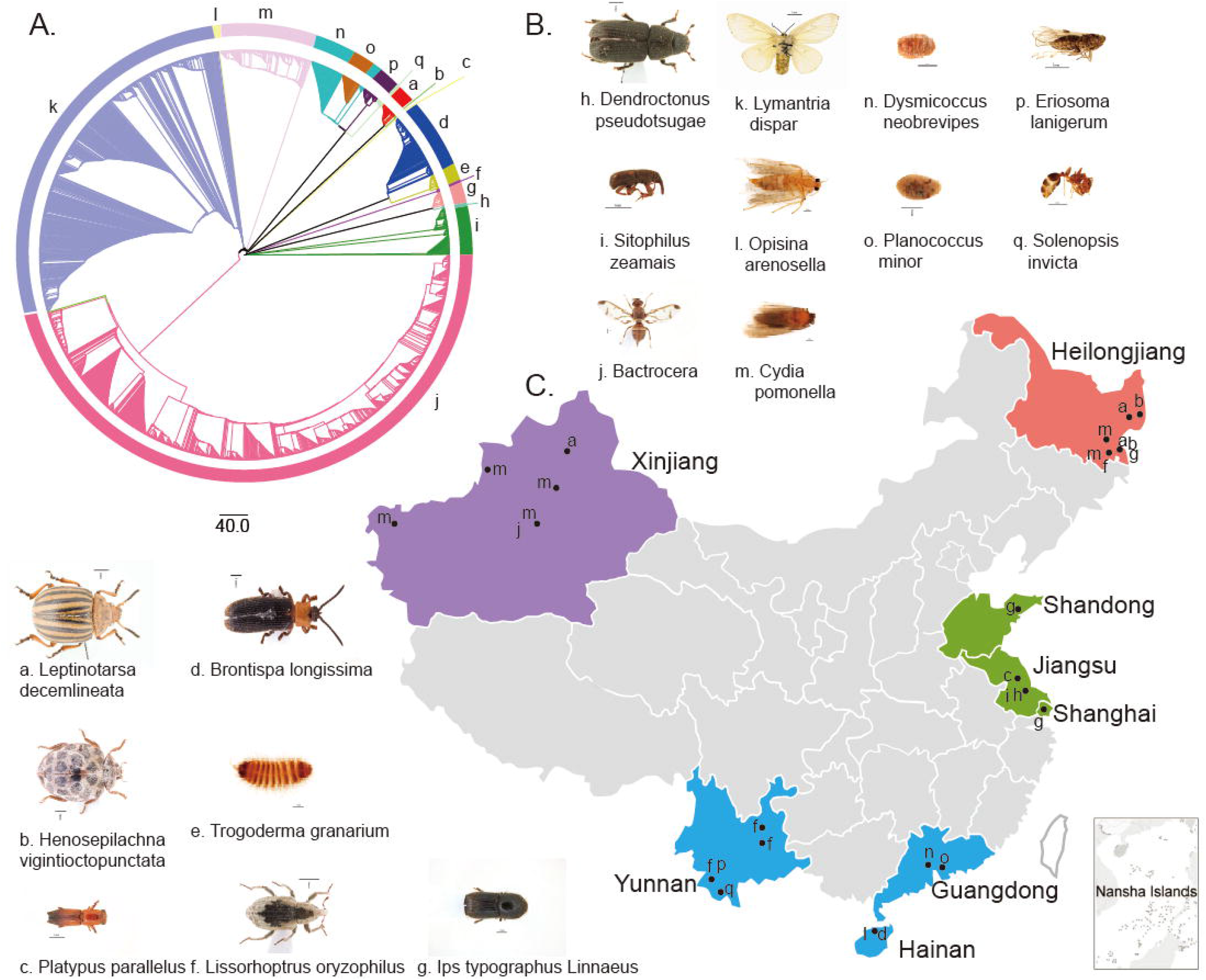
Collection of insect samples and analysis of evolutionary relationship. **A**. Phylogenetic tree of insects using the COI gene. Each color represents a different species. The lower-case letters represent species that correspond to those in the photo and map below. **B**.Photos of sampled insects. **C**. Insect sampling sites. The sampling sites are indicated on the map of China, and the four main collection areas are marked with four different colors. J is *Carpomya vesuviana* Costa.

The tested insects varied greatly in size. To ensure that gene extraction was not affected by insect size, all insect species were divided into three groups according to their body length. If the body length of the adult insect was >5 mm, such as in *Leptinotarsa decemlineata, Ips typographus*, and *Lymantria dispar*, one individual was used as one sample for gut microbiota analysis. If the body length ranged from 2 to 5 mm, such as in *L. oryzophilus, C. pomonella*, and *Bactrocera cucurbitae*, five individuals collected from one site were used as one sample for DNA extraction. If the body length was < 2 mm, such as in *Trogoderma granarium, Eriosoma lanigerum*, and *Planococcus minor*, 10 insects collected from one site were used as one sample to extract DNA. The purpose of several individuals making up a sample was to more effectively extract DNA and also to reduce individual differences and human impact. A total of 179 gut microbiota samples from quarantine insects were obtained and subjected to 16S rDNA sequencing (Table S1).

### Sequencing and analysis of microbiota diversity

All 179 samples were simultaneously sequenced, producing a total of 16,465,986 reads. Each sample contained 50,000–150,000 reads and produced around 15-37 Mbp of data (**Figure 2A**). All data were submitted to the Genome Sequence Archive (GSA), and the accession ID is CRA002386. After quality control, there were 50,000–125,000 reads with 15–32 Mbp remaining for each sample (**Figure 2A**). Clean data were then classified by the QIIME software [12]. We classified 1527 taxa of microbes belonging to two kingdoms: Archaea and Bacteria. There were 38 phyla, 98 classes, 178 orders, 277 families, and 598 genera (**Figure 2B**). All 1527 taxa were identified to various taxonomic levels. All 1527 could be identified to the kingdom level, 1525 to the phylum level, 1505 to the class, 1450 to order, 1320 to family, 989 to genus, and 290 to the species level (**Figure 2B**).

**Figure 2.**
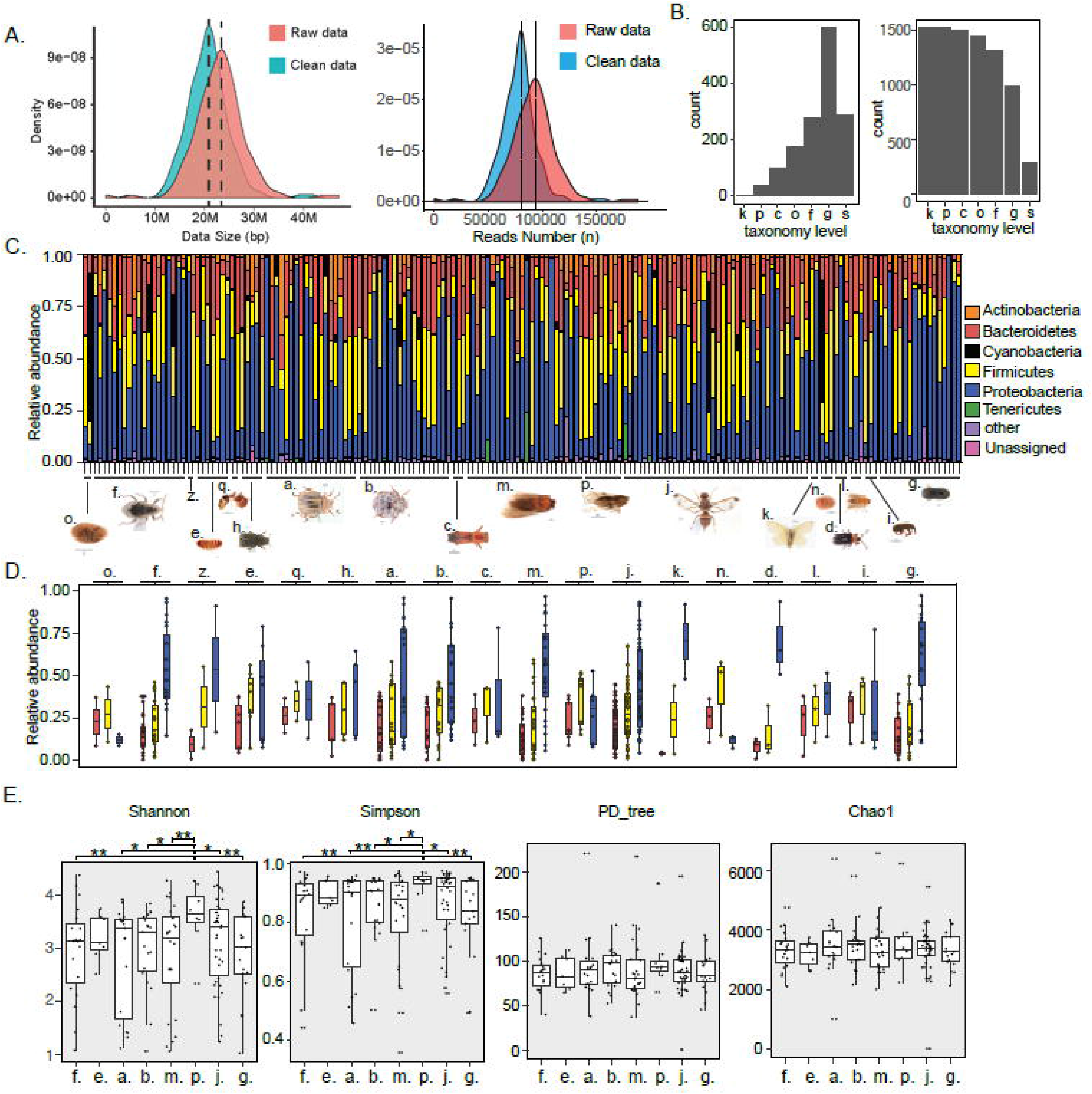
Density distribution of sequencing quantity. **A**. The left panel is the data size in base pairs (bp) per sample (M = million). The right panel is the sequence number per sample. **B**. The left panel is the number of different taxa identified at each taxonomic level. The right panel is the number of microbes that could be identified at each taxonomic level (k = kingdom, p = phylum, c = class, o=order, f = family, g = genus, s = species). **C**. Compositions of gut microbiota for each sample at the phylum level. **D**. Relative abundance of three dominant phyla in each insect species. The color of the box corresponds to that in C. **E**. Alpha diversity of eight representative insect species (**, P < 0.01. *, P < 0.05).

Due to the complexity of the data, we analyzed the composition of the microbiota for each sample at the phylum level. Bacteroidetes, Firmicutes, and Proteobacteria were the three dominant phyla in each sample. Actinobacteria were common and ubiquitous (**Figure 2C**). Cyanobacteria and Tenericutes were observed in some samples. The proportions of these phyla were different in each sample due to the different host species and different collection sites. Of the three dominant phyla, Proteobacteria had the maximum proportion, followed by Firmicutes and Bacteroidetes (**Figure 2D**). Insect species whose sample number was >5 were subjected to alpha diversity comparison. Except for *Eriosoma lanigerum*, all other species exhibited no significant difference in either Shannon, Simpson, PD_tree or the Chao1 index (**Figure 2E**). However, the Shannon index and Simpson index of *Eriosoma lanigerum* (p. in **Figure 2E**) showed significant differences compared to the other insects except *Trogoderma granarium* (e. in **Figure 2E**).

### Screening of methods for determining geographical source based on microbiota

To link the geographic source and microbiota of quarantine insects, we analyzed microbiota data of all insects collected from the field using principal component analysis (PCA). However, the dots representing insects from five geographical areas were tangled, as shown in **Figure 3A** and Figure S1. In a previous study, multiple factors were shown to affect the gut microbiota of insects. These included insect species, developmental stage, diet, sex, and geographical location [13–14]. According to the ADONIS test results based on normalized abundance of gut microbiota, the effect size of geographical factor (R^2^=0.016) was a little higher than sex factor (R^2^=0.007), but lower than effect of developmental stage (R^2^=0.028) and insect species (R^2^=0.09). This suggested that the geographical source of the insect was not the dominant factor affecting the gut microbiota. Therefore, we used linear discriminant analysis (LDA), a classification algorithm based on prior information. We compared PCA and LDA by ADONIS test using reared insects whose geographical source and diet factor were controlled. Given sex as prior information, LDA (R^2^=0.67, P-value=0.001) was better able to discriminate sex compared to PCA (R^2^=0.01, P-value=0.881) (**Figure 3B**, Figures S2Aand S3A). For discriminate insects species, a similar conclusion (R^2^=0.71, P-value=0.001 for LDA, R^2^=0.10, P-value=0.36 for PCA) applied when species information was given to the LDA (Figures S2B, and S3B). These results suggested that LDA has a better diagnostic ability than PCA to extract specific factors affecting gut microbiota (**Figures 3E**).

**Figure 3.**
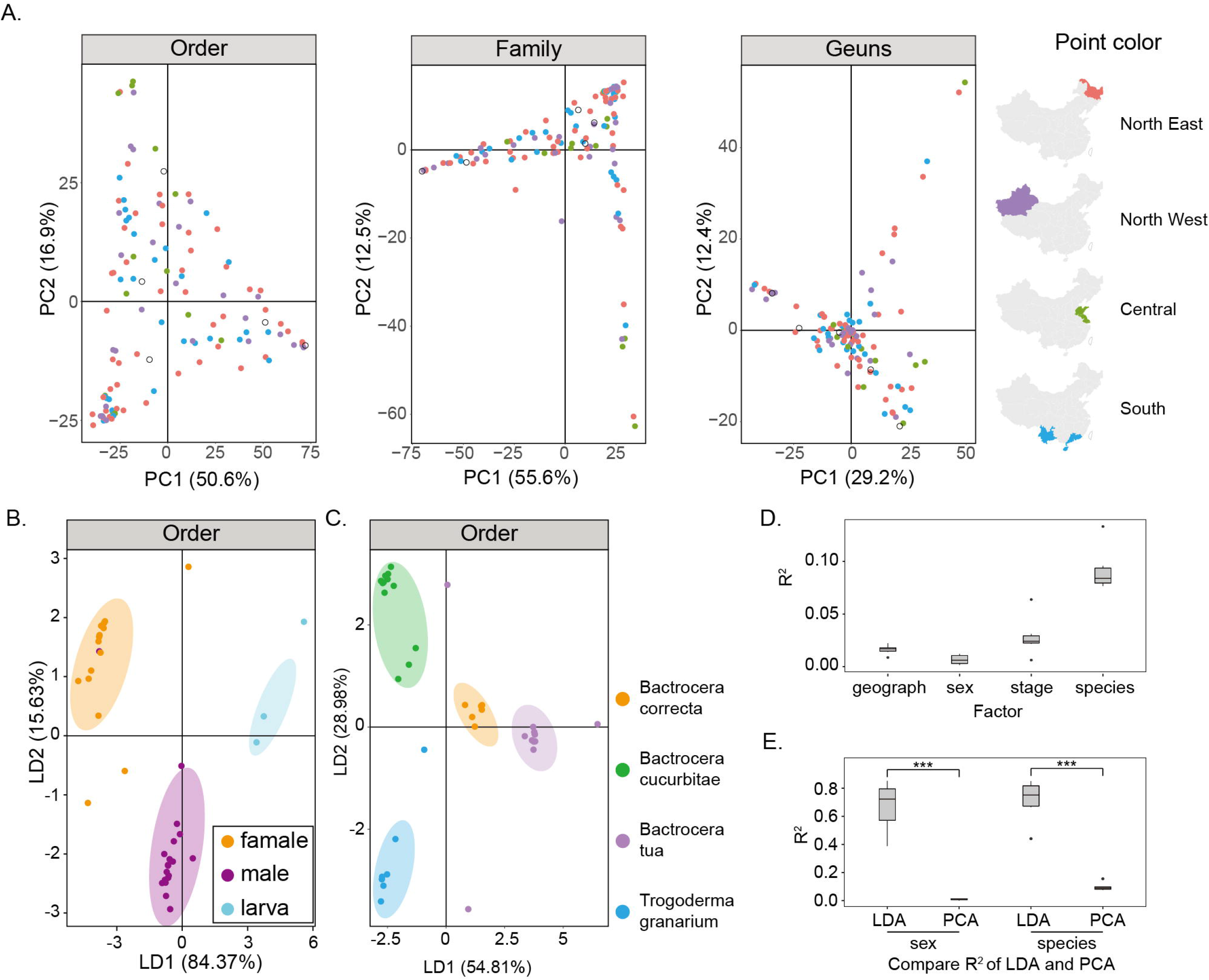
Screening of methods for determining geographical source based on insects’ microbiotaA. PCA method of distinguishing insects collected from five geographical areas. Result of principal component analysis (PCA) showed that dots representing insects from five geographical regions were intertwined and difficult to distinguish. **B**. LDA methods of distinguishing insects considering the sex factor. Result of linear discriminant analysis (LDA) showed that dots can gather into clusters, which can distinguish insects of different sexes. **C**. LDA methods of distinguishing insects considering the insect species factor.These dots can gather into clusters, which can distinguish insects of different species.. D. Effect size of each insect related factor. (R2 was calculated from ADONIS test). E. Comparison of ADONIS effect size between LDA and PCA using t-test. (**, P<0.01; ***, P<0.001)

### Verification of LDA’s ability to distinguish geographical source

To determine the ability of LDA to find the taxonomic level of microbes given geographical information, we performed LDA at all taxonomic levels of microbes except for kingdom. Insects from the same geographical area (marked by dots in **Figure 4A)** were grouped together at the class, order, family, and species levels. However, the insects from different geographical areas could not be distinguished at the phylum or genus levels of the microbes (**Figure 4A**). The dots representing insects collected abroad were located far from the dots representing insects collected in China. To determine the robustness of this method, a Jackknife was performed 1000 times at all taxonomic levels of microbes except for the kingdom level by dropping approximately 15% of the sample each time. The distribution of accuracy from the jackknife method showed that accuracy at the phylum level was lowest, while class level and order level were higher than 0.95 (**Figure 4B**). Bootstrapping was also performed 1000 times, and a similar result was obtained (**Figure 4C**). Class and order were considered as proper taxonomic levels to distinguish geographical clusters of host insects because of the high accuracy and low standard deviation in both jackknife and bootstrap techniques (**Figure 4D and E**).

**Figure 4.**
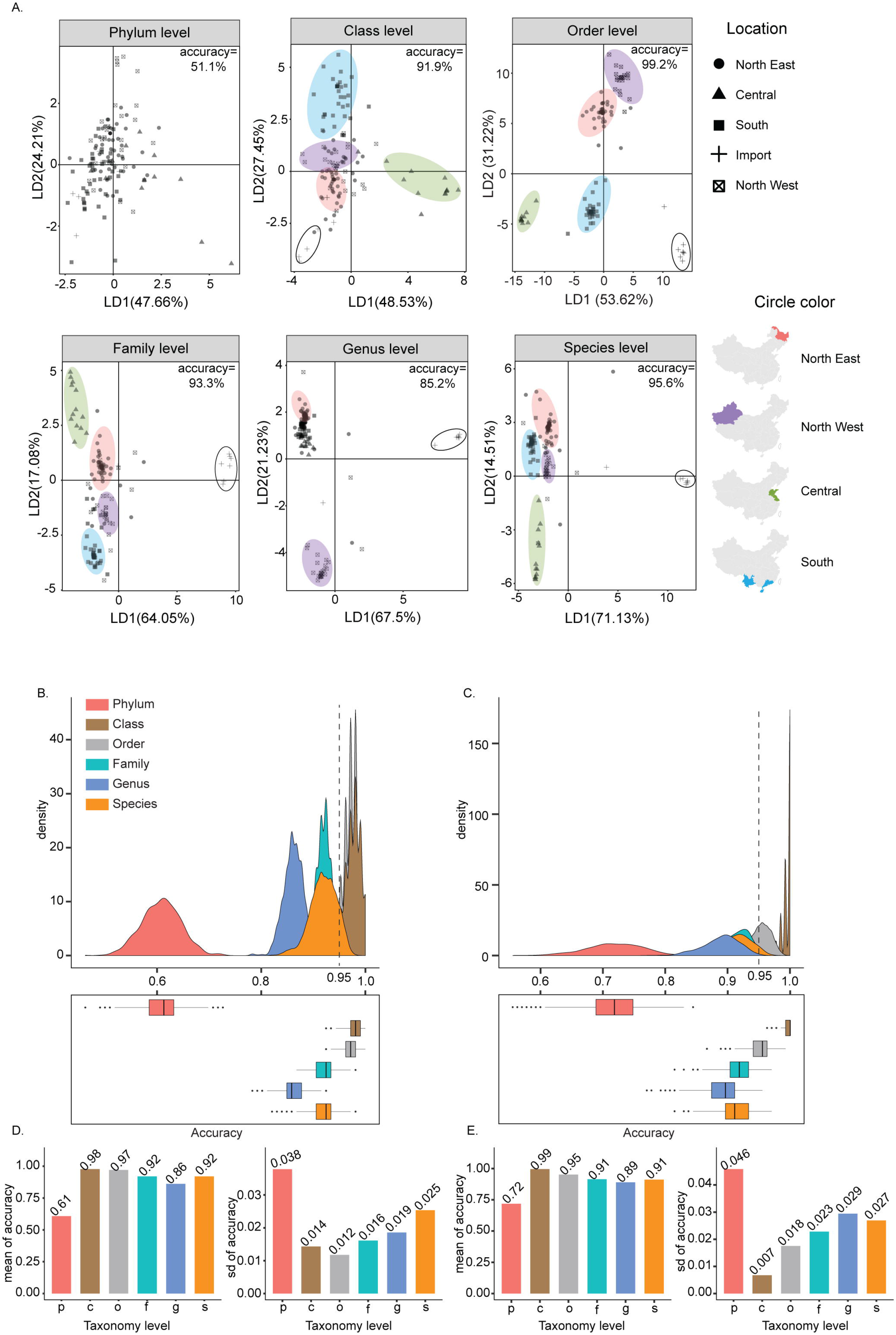
Verification of LDA’s ability to distinguish all insects at each microbial taxonomic level. **A**. The points in the linear discriminant analysis (LDA) were shaped and circled by sampling the geographic area. The transparent circle delineates several insects that were intercepted at entry ports. The dots representing insects collected abroad were located far from the dots representing insects collected in China. Class and order were considered as proper taxonomic levels to distinguish geographical clusters of host insects. **B**. Distribution of accuracy at each taxonomic level of gut microbes via the jackknife. **C**. Distribution of accuracy at each taxonomic level of gut microbes via the bootstrap. **D**., **E**. Mean and SD of accuracy for the jackknife and bootstrap. The high accuracy and low standard deviation were verified with the jackknife and bootstrap.

### LDA distinguishing insects from different geographical source

To study the relationship between gut microbiota and host geographical source, the factor of insect species was controlled, and LDA using geographical information was performed using five insect species. These were *Cydia pomonella, Ips typographus, Leptinotarsa decemlineata, Lissorhoptrus oryzophilus*, and *Henosepilachna vigintioctopuntata*, each of which was sampled from at least three areas. The *Cydia pomonella* result is shown in **Figure 5**, and the results were similar for the other four selected insects (Figures S4 and S5). A total of 18 *Cydia pomonella* were collected from four sites. Two sites, Dongning and Mudanjiang, are in the northeast of China, while Urumqi and Korla are located in the northwest. In **Figure 5**, using the ADONIS test (Figure S6 A and B), those four sampling sites could be distinguished at most taxonomic levels of microbes (P-value=0.04). There was not sufficient discrimination (P-value=0.32) at the genus level when we analyzed the gut microbiota of *Henosepilachna vigintioctopuntata* (Figures S5B and S6F), and discrimination at the species level was inadequate (P-value=0.33) in the analysis of the distribution of *Lissorhoptrus oryzophilus* (Figures S5A and S6E). Class level (P-value=0.04), order level (P-value=0.04), and family level (P-value=0.06) information were available, but the phylum level (P-value=0.009) showed better discriminatory ability (Figures S4, S5, and S6A-G). The phylum level was the best for tracing a single insect species (Figure S6G).

**Figure 5.**
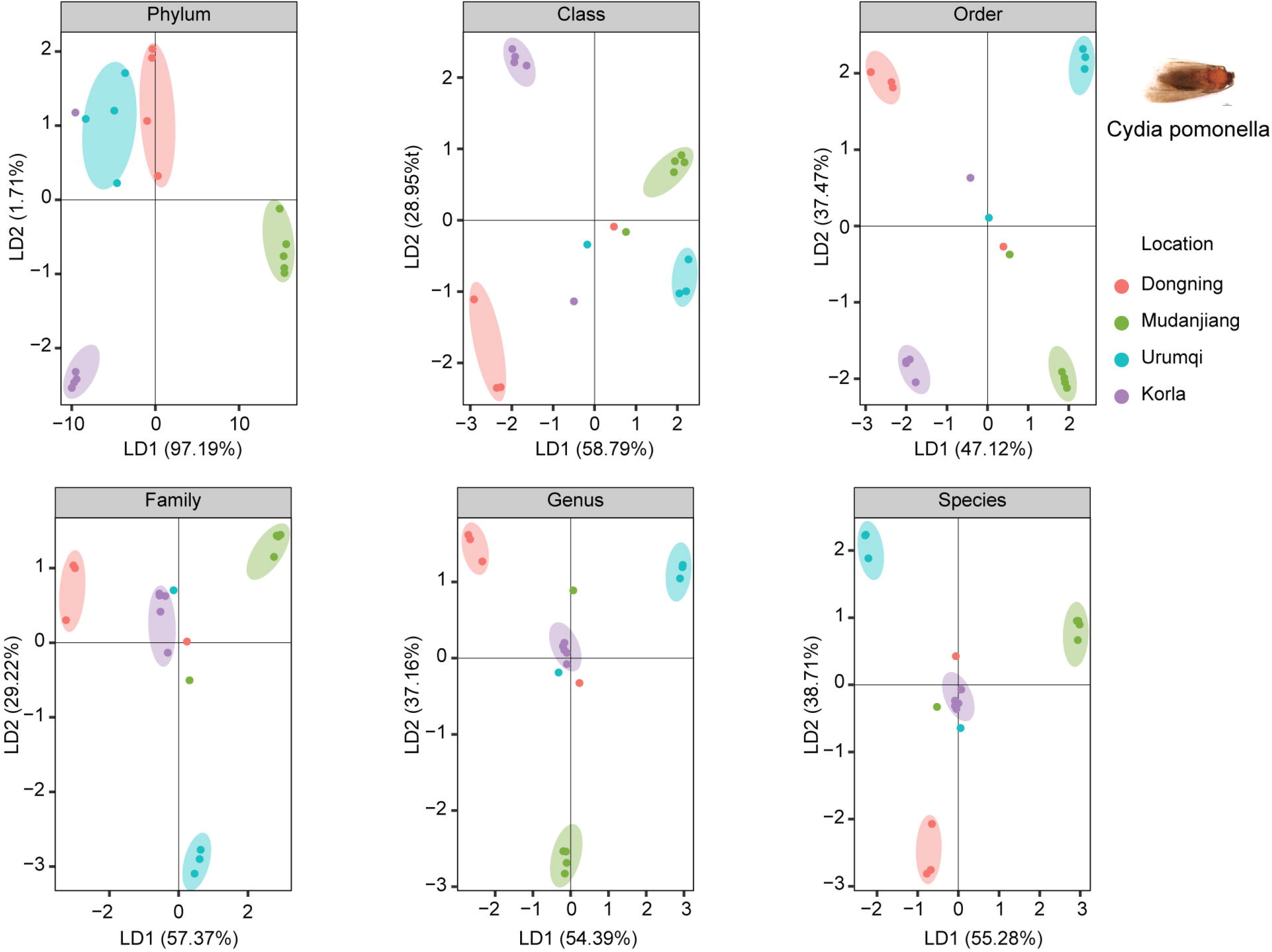
Application of LDA method in each taxonomic level of gut microbes of *Cydia pomonella*. A total of 18 *Cydia pomonella* were collected from four sites. Two sites, Dongning (red dots) and Mudanjiang (green dots), are in the northeast of China, while Urumqi (blue dots) and Korla (purple dots) are located in the northwest. Dots of the same color were distributed in clusters, showing that four sampling sites could be distinguished at most taxonomic levels of microbes.

Each of the five insects mentioned above was analyzed at the phylum level of gut microbiota. *Cydia pomonella* (P-value=0.01, R^2^=0.88, **Figure 6A**, Figure S6H), *Ips typographus* (P-value=0.004, R^2^=0.86, **Figure 6B**, Figure S6I), and *Leptinotarsa decemlineata* (P-value=0.002, R^2^=0.86, **Figure 6C**, Figure S6J) could be traced accurately. For *Lissorhoptrus oryzophilus*, there was an overlap between the Xundian group and the Menglian group (P-value=0.002, R^2^=0.59, **Figure 6D and** Figure S6K). Both of these species were located in Yunnan province. The Suifenhe group overlapped with the Hulin group for *Henosepilachna vigintioctopuntata* (P-value=0.001, R^2^=0.63, **Figure 6E) (**Figure S6L), though it is closer to Dongning in geography. A heat map was used to find the features of the microbe phylum with the closest relation to the host geographical source using relative abundance. A geographical correlation was not obvious using one or several microbial phyla, but there was some evidence for a pattern (**Figure 6**).

**Figure 6.**
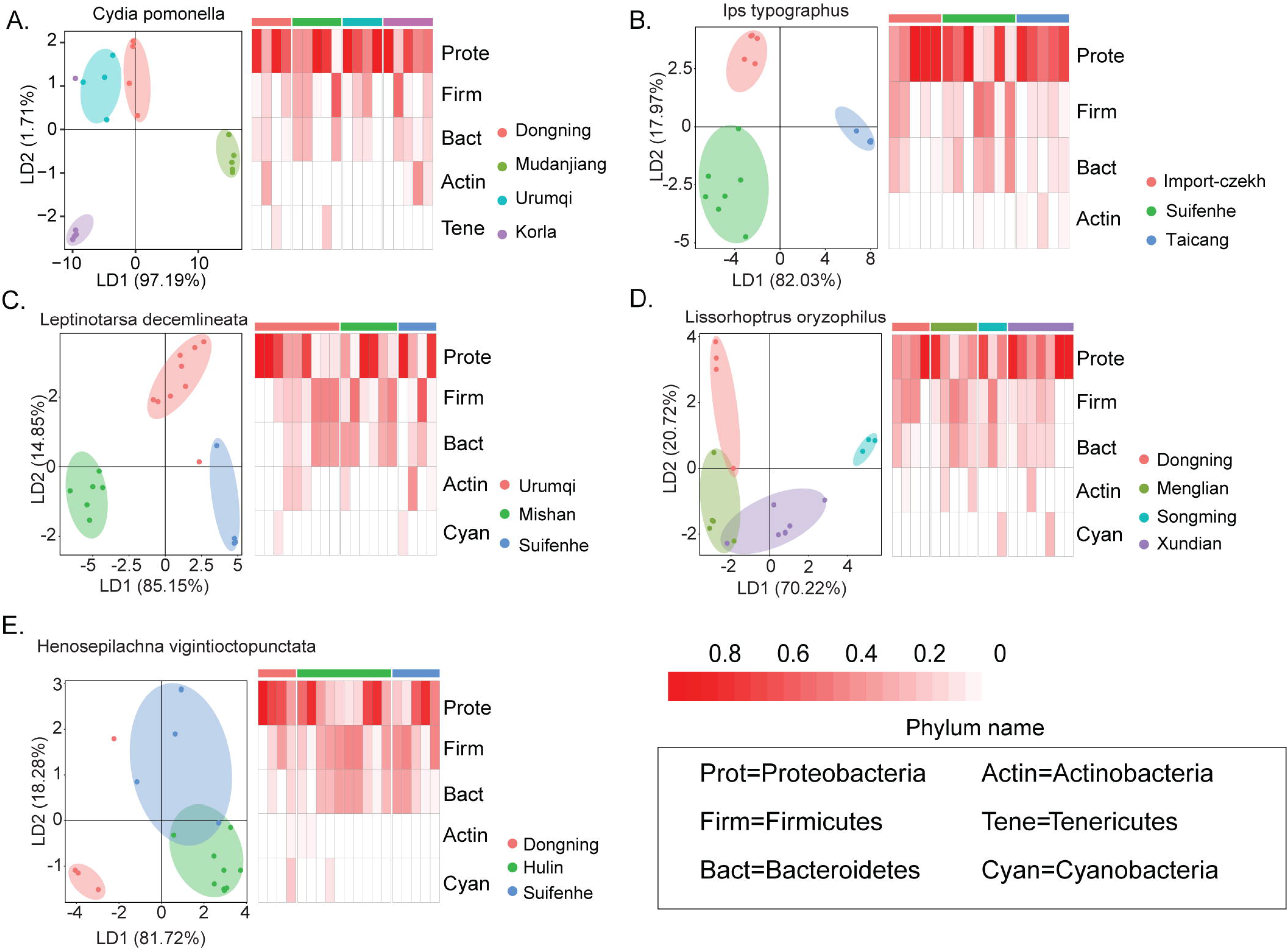
LDA and heatmap for gut microbes of five representative insects. Heatmap showing relative abundance of each insect at the phylum level. The bar on top of the heatmap represents the geographical source of the insect. Each of the five insects was analyzed at the phylum level of gut microbiota. *Cydia pomonella* (A), *Ips typographus* (B), and *Leptinotarsa decemlineata* (C) could be traced accurately. For *Lissorhoptrus oryzophilus* (D), there was an overlap between the Xundian group and the Menglian group. Both of these species were located in Yunnan province. The Suifenhe group overlapped with the Hulin group for *Henosepilachna vigintioctopuntata* (E), though it is closer to Dongning in geography.

## Discussion

To the best of our knowledge, this is the first attempt to trace quarantine insects based on gut microbiota. There is no effective and fast method currently available to identify the geographical source of quarantine insects. Although the methods of DNA barcoding and RFLP can predict the geographical origin, it takes many insect generations to detect gene flow from multiple origins [15,16]. These methods are better at identifying the genetics of evolutionary relationships or for working with species complexes [17,18].

Several methods were evaluated in this study. The method represented by PCA may not be the most appropriate approach to extracting geographical information. This is because species, sex, developmental stage, diet, and geographical source of the host all affect the gut microbiota, while the geographical source of the insects may not be the principal factor. We found that methods based on prior information such as LDA were better for distinguishing the geographical sources of insects using the gut microbiota. The geographical factor was revealed by the composition patterns of gut microbes. Supervised machine learning is effective at extracting composition patterns from data using prior information to establish a prediction model. Therefore, supervised learning is our recommended strategy. Some outliers were observed in the LDA analysis because of the prior information. The sampling sites of insects were defined as the geographical sources. However, the insects might have originated in other places, and their gut microbiota may not have been localized. This situation is inevitable in sampling, but an increased number of samples and a robust algorithm may help to reduce this noise.

This study showed that different insect species can be distinguished using the phylum level of gut microbiota. A single insect species from different regions can be distinguished at the genus level of microbiota. For invasive species in China with limited distributions such as *C. pomonella*, this method can easily and quickly identify the source of the invasion. Similarly, for widely distributed species like *I. typographus*, this method can distinguish specimens from China and abroad. This is a problem that cannot be addressed by genetic tests or morphology. The insect gut microbiota is closely related to host states of sex, diet, developmental stage, and niche occupation. For example, honeybees living in different locations can differ significantly in their gut microbiota. The structure of the microbial community can also differ among bees depending on whether or not they are foraging on flowering rape crops [19]. Worker honeybees and solitary bees also have different gut microbiota. There are eight distinct bacterial species or phylotypes in worker honeybees, three of which are Gram-positive species, for example *Bifidobacterium*, and five Gram-negative species includingβ-proteobacterium [20]. In predatory insects, the diversity of prey consumed can increase the diversity of bacteria in the gut [21]. Meanwhile, the diversity of microorganisms is also greatly influenced by differences in plants consumed [22–24]. The gut community of *L. decemlineata* larvae feeding on tomato was dominated by the genera *Stenotrophomonas* and *Lactococcus*, and larvae feeding on potato mainly had *Enterobacter* [25].

The accuracy of 16S amplifier sequencing was affected by the depth of sequencing and the depth of coverage. The sequencing results showed that differences in gut microbiota at the phylum level can distinguish insect species, and differences at the genus level can distinguish the same insect species from different source locations. It is feasible to trace the origin of insects from different geographic sources using the genus level of gut microbes. However, due to the limited number of sequencing samples, the LDA values of some samples did not cluster. These included *C. pomonella* collected from Ili and *L. decemlineata* from Urumqi. However, for *C. pomonella* and *I. typographus*, their native individuals and intercepted conspecifics were distinguishable in this study.

This paper also addressed the problem of determining the gut microbiota in small insects. The body length of different insect species varies greatly. Most studies of insect gut microbes have dissected the gut and evaluated the interaction between microbiota and host stage for bacteria identification [26–30]. However, it is difficult to obtain individual guts from very small insects. Some individuals represented one sample from which the genomic DNA was extracted, and the microbiota was amplified from the entire insect body. This approach may help solve the problem of individual differences and small samples. The methods of genomic DNA extraction from insect gut microbiota need further study to establish better standards for future research.

In summary, we developed a method to trace the origin of quarantine insects using a prior information based LDA. Bacteroidetes, Firmicutes, and Proteobacteria were the three dominant phyla in insect guts, and their relative abundance differed among insect species. Class and order level of gut microbiota can provide geographic information even though the gut microbiota is masked by insect species differences. For a single insect species, the class, order, and family levels of gut microbiota are useful taxonomic levels. Although this new method can determine the geographical source of many insects without requiring species identification, its popularization and application must be based on a database of insect gut microorganisms. If the insect species can be identified, the geographical source of the insect might be more accurately located. The quantity and quality of basic data will have a large impact on identification accuracy, especially for prior information-based methods. The gut microbiota of insects is complex and variable. A high-quality background database could improve the accuracy and stability of the identification model. To establish this database, multiple insect species should be sampled, and the samples should be taken from different geographical areas. To control the batch effects and ensure database quality, the sample processing methods should be consistent.

## Materials and methods

### Sample preparation

Quarantine insects in China were used for this study. The collection sites were determined based on previous monitoring records of quarantine insects. Most insects were captured with a net. At each location, we collected at least 10 insect individuals for each species. Each insect was alive before it was immersed into RNAlater® Stabilization Solution (Cat. No. AM7021, Ambion, Austin, Texas, USA).

### DNA extraction, 16S rRNA gene amplification and sequencing

We used three gut microbiota DNA extraction strategies based on the insect body length. Large individuals (> 5 mm) were extracted as one sample. Five intermediate size individuals (2–5 mm) or ten small individuals (< 5 mm) were extracted as one sample. Before extraction, the surface of each insect was sterilized with 70% ethanol and washed twice with sterile PBS [31]. The insect abdomens or the whole insects were put into a special EP tube weighing 0.3 g along with 0.1 mm glass beads (Cat. No. 11079101, BioSpec, Oklahoma City, OK, USA). DNA extraction was done according to the QIAamp® Fast DNA Stool Mini Kit (Cat. No. 51604, Qiagen, Hilden, Germany). Samples were pretreated before the DNA extraction. Pretreatment steps were as follows: 1.4 ml of inhibit EX buffer was first added into the EP tube and the sample was ground in a bead beater for 1 min. Second, the samples were incubated at 95°C for 10 min, and then reground for 2 min. The samples were then centrifuged for 1 min to pelletize the particles. The following steps were the same as those given in the kit protocol. The concentrations of DNA genes were measured with Nanodrop, using this as a template for PCR amplification.

The PCR reaction system used the protocol of HiFi HotStart DNA Polymerase (Cat. No. KR0369, Kapa Biosystems, Boston, MA, USA), and the experiments were carried out based on a two-step PCR reaction. The first PCR amplification was performed with the primer pairs under the following conditions: a denaturing step at 95°C for 5 min followed by 20 cycles of 98°C for 20 s, 52°C for 30 s, 72°C for 30 s, and a final step of 5 min at 72°C. The primer pairs were 5’-CCTACGGGNBGCASCAG-3’ (forward) and 5’-GACTACNVGGGTATCTAATCC-3’ (reverse). The PCR products were purified with the kit from Agencourt AMPuer XPsystem (Cat. No. A63880, Beckman Coulter, South San Francisco, CA, USA), and the purified products were used with the second PCR amplification. The second PCR amplification conditions were a denaturing step at 95°C for 5 min followed by 10 cycles of 98°C for 20 s, 60°C for 30 s, 72°C for 30 s, and a final step of 5 min at 72°C. These primer pairs were the Illumina sequencing joint with different index, having the V3 and V4 information of 16S. The PCR products were purified using the same protocol as above. Then, the concentrations of each sample were detected after electrophoresis. The method of sequencing was paired-end 250 bp (PE250) sequencing (HiSeq2500, Illumina, San Diego, CA, USA).

### Quality control and taxonomy assignment

After sequencing, sequences were distributed into samples based on barcodes. After we removed barcodes and primers, we trimmed 10 bp at the start and end of each read for quality control [32]. Sequences longer than 104 bp were retained after trimming bases whose quality was below 20 using Sickle V1.33 software. Error correction was performed by SPAdes V3.1.9 software [33]. The workflow pick_open_reference_otus.py in QIIME v1.9.1 was used to pick OTUs at 97% similarity and to assign a species level using the UCLUST method in the GREEN GENE database.

### Phylogenetics

We downloaded the COI gene sequences of the test insects from the Barcode of Life Data System (BOLD) database [34]. These 2374 sequences were aligned using Clustal Omega (https://www.ebi.ac.uk/Tools/msa/clustalo/). Then, a method based on maximum likelihood, RAxML-HPC2 v8.2.10 [35], was used to construct a phylogenetic tree. We performed 1000 bootstrap replicates for this tree after removing suspicious sequences. The tree was edited and visualized by Figtree v1.4.3.

### Data analysis and visualization

The density distribution of sequencing quantity and composition of gut microbiota were analyzed using R v3.4.1. The Alpha diversity index was calculated using vegan package v2.5-3 in R. Principal component analysis (PCA) was performed using R stats package v3.4.1. Linear discriminant analysis (LDA) was performed using the MASS package in R software. The heatmap was visualized using the R package pheatmap v1.0.10.

PCA for insects collected from five geographical areas was performed based on relative abundance of microbes. LDA for all insects that were collected from five geographical areas was performed based on relative abundance normalized using the log function. LDA for reared insects and representative insects was performed using relative abundance. The first and second components were chosen for visualization using the R package ggplot2. The ratio of the LDA classifying results to the original sample information was defined as the accuracy. A Jackknife was performed 1000 times, excluding 15% of the sample from each geographical source each time. Bootstrap resampling was performed 1000 times, and the number of bootstrap samples was equal to the original samples. A heatmap was constructed using the relative abundance at the order level using the R package pheatmap. Distance between each group in LDA or PCA was measured with a permutational multivariate analysis of variance (PERMANOVA) in the R package vegan v2.5-3. Comparison of ADONIS R2 and P value between LDA and PCA was performed with t-test by R package vegan v-2.5-3.

### Availability

All of the sequencing data have been deposited in Genome Sequence Archive (GSA: CRA002386), and the data can be available freely at https://bigd.big.ac.cn/search?dbId=gsa&q=CRA002386.

## CRediT author statement

Yanxue Yu: Resources, Data curation, Writing-Original draft preparation. Qi Wang: Formal analysis, Writing-Original draft preparation. Ping Zhou: Investigation. Na Lv:Methodology. Wei Li:Software. Fangqing Zhao: Conceptualization, Methodology. Shuifang Zhu:Supervision. Di Liu: Conceptualization, Writing-Review&Editing.

## Competing interests

The authors have declared no competing interests.

## Acknowledgments

This work was supported by grants from the National Key Research and Development Program of China (Grant Nos. 2016YFC1200800, 2016YFC1200803, 2016YFC1200805, 2018YFC0809100). ACCDON LLC language editing improved the wording of this report.

## Supplementary material

**Table S1 Detailed information of insect samples**

**Figure S1 Distinguishing ability of PCA for tracing the geographical source of insects**

Each point represents an insect, and the point color represents the geographical area where the insect was collected. Hollow points represent the imported insects.

**Figure S2 LDA for discriminating sex or species of insects based on gut microbes**.

The linear discriminant analysis (LDA) can distinguish insects with sex (**A**) and species differences (**B**).**Figure S3 Discrimination comparison of LDA and PCA using the PERMANOVA test**

The methods of principal component analysis (PCA, **left panel**) and linear discriminant analysis (LDA, **right panel**) were compared by ADONIS test using reared insects whose geographical source and diet factor were controlled. **A**. Comparison of two methods in sex factors. LDA was better able to discriminate sex compared to PCA. **B**. Comparison of two methods in species factors. For discriminate insects species, R^2^=0.71, p-value=0.001 for LDA, R^2^=0.10, p-value=0.36 for PCA. These results suggested that LDA has a better diagnostic ability than PCA to extract specific factors affecting gut microbiota.

**Figure S4 Verification of LDA’s ability to distinguish *Ips typographus* and *Leptinotarsa decemlineata* at each microbial taxonomic level**

**A**. The linear discriminant analysis (LDA) distinguished *Ips typographus* at all taxonomic level. A point represented an insect, the point color showed its geographical source, and the same geographical area was clustered together. Red areas indicated samples from abroad, green areas indicated samples from Suifenhe in Heilongjiang Province, and blue areas indicated samples from Taicang in Jiangsu Province. **B**. The linear discriminant analysis (LDA) distinguished *Leptinotarsa decemlineata* at all taxonomic level. A point represents represented an insect, and the point color shows showed its geographical source, and the same geographical area was clustered together. Red areas indicated samples from Urumqi, green areas and blue areas respectively indicated samples from Mishan and Suifenhe in Heilongjiang Province..

**Figure S5 Verification of LDA’s ability to distinguish *Lissorhoptrus oryzophilus* and *Henosepilachna vigintioctopuntata* at each microbial taxonomic level**. A point represents an insect, and the point color shows its geographical source. **A**. The linear discriminant analysis (LDA) distinguished *Lissorhoptrus oryzophilus* at all taxonomic level. A point represented an insect, the point color showed its geographical source, and the same geographical area was clustered together. Red areas indicated samples from Dongning in Heilongjiang Province, green areas indicated samples from Menglian, and blue areas indicated samples from Songming, and purple areas indicated samples from Xundian in Yunnan Province. **B**. The linear discriminant analysis (LDA) distinguished *Henosepilachna vigintioctopuntata* at all taxonomic level. A point represents represented an insect, and the point color shows showed its geographical source, and the same geographical area was clustered together. Red areas indicated samples from Dongning, green areas and blue areas respectively indicated samples from Hulin and Suifenhe in Heilongjiang Province.

**Figure S6 PERMANOVA test for five insect species**

P values of PERMANOVA tests at each taxonomic level of gut microbiota for *Cydia pomonella* **(A, B)**, *Ips typographus* **(C)**, *Leptinotarsa decemlineata* **(D)**, *Lissorhoptrus oryzophilus* **(E)**, and *Henosepilachna vigintioctopuntata* **(F) G**. The P values of all five insect species. PERMANOVA test of distance between each group in LDA for *Cydia pomonella* **(H)**, *Ips typographus* **(I)**, *Leptinotarsa decemlineata* **(J)**, *Lissorhoptrus oryzophilus* **(K)**, and *Henosepilachna vigintioctopuntata* **(L)**.

