## Supplemental Table 1 for "First Glimpse of Gut Microbiota of Quarantine Insects in China"

**Table 1 Detail information of insect samples**

| **Sample ID** | **Scientific name** | **Collection Sites** | **DNA Concentration (ng/μl)** | **Raw data(mb)** | **Clean data(mb)** | **Raw reads** | **Clean reads** |
| --- | --- | --- | --- | --- | --- | --- | --- |
| 205 | Planococcus minor | Guangzhou of Guangdong Province | 23.68 | 22 | 19 | 69454 | 59587 |
| 206 | Planococcus minor | Guangzhou of Guangdong Province | 64.23 | 32 | 28 | 102550 | 89754 |
| 36 | Lissorhoptrus oryzophilus | Dongning of Heilongjiang Province | 22.22 | 26 | 24 | 87298 | 76426 |
| 37 | Lissorhoptrus oryzophilus | Dongning of Heilongjiang Province | 22.13 | 30 | 25 | 94464 | 81592 |
| 38 | Lissorhoptrus oryzophilus | Dongning of Heilongjiang Province | 14.68 | 24 | 21 | 78560 | 69385 |
| 39 | Lissorhoptrus oryzophilus | Dongning of Heilongjiang Province | 11.01 | 18.6 | 17 | 61918 | 51086 |
| 122 | Lissorhoptrus oryzophilus | Menglian of Yunnan Province | 5.84 | 26 | 22 | 80860 | 67087 |
| 123 | Lissorhoptrus oryzophilus | Menglian of Yunnan Province | 5.96 | 30 | 26 | 99824 | 77654 |
| 190 | Lissorhoptrus oryzophilus | Menglian of Yunnan Province | 3.87 | 28 | 25 | 91726 | 79422 |
| 191 | Lissorhoptrus oryzophilus | Menglian of Yunnan Province | 2.84 | 26 | 22 | 81492 | 70433 |
| 192 | Lissorhoptrus oryzophilus | Menglian of Yunnan Province | 8.63 | 26 | 24 | 86096 | 73955 |
| 193 | Lissorhoptrus oryzophilus | Songming of Yunnan Province | 4.73 | 30 | 26 | 94114 | 81426 |
| 194 | Lissorhoptrus oryzophilus | Songming of Yunnan Province | 4.8 | 24 | 21 | 77630 | 67053 |
| 195 | Lissorhoptrus oryzophilus | Songming of Yunnan Province | 5.18 | 28 | 25 | 91036 | 79774 |
| 53 | Lissorhoptrus oryzophilus | Xundian of Yunnan Province | 17.11 | 36 | 31 | 116608 | 98949 |
| 54 | Lissorhoptrus oryzophilus | Xundian of Yunnan Province | 12.06 | 38 | 33 | 124876 | 102255 |
| 55 | Lissorhoptrus oryzophilus | Xundian of Yunnan Province | 5.35 | 24 | 20 | 77240 | 60928 |
| 153 | Lissorhoptrus oryzophilus | Xundian of Yunnan Province | 1.45 | 32 | 29 | 105502 | 89092 |
| 154 | Lissorhoptrus oryzophilus | Xundian of Yunnan Province | 1.13 | 26 | 24 | 86758 | 74504 |
| 155 | Lissorhoptrus oryzophilus | Xundian of Yunnan Province | 3.75 | 26 | 23 | 85622 | 72870 |
| 156 | Lissorhoptrus oryzophilus | Xundian of Yunnan Province | 1.56 | 18.6 | 17 | 62034 | 54202 |
| 161 | Bactrocera correcta | Beijing Lab rearing | 6.86 | 32 | 28 | 104232 | 92355 |
| 162 | Bactrocera correcta | Beijing Lab rearing | 7.14 | 36 | 32 | 117478 | 99571 |
| 163 | Bactrocera correcta | Beijing Lab rearing | 73.39 | 24 | 20 | 73674 | 63719 |
| 164 | Bactrocera correcta | Beijing Lab rearing | 9.35 | 19.8 | 18 | 66182 | 57700 |
| 199 | Bactrocera correcta | Beijing Lab rearing | 12.48 | 28 | 24 | 89756 | 70705 |
| 200 | Bactrocera correcta | Beijing Lab rearing | 12.4 | 24 | 22 | 80138 | 70080 |
| 207 | Phenacoccus solenopsis | Guangzhou of Guangdong Province | 25.36 | 26 | 22 | 82318 | 71654 |
| 208 | Phenacoccus solenopsis | Guangzhou of Guangdong Province | 37.4 | 28 | 24 | 88752 | 77993 |
| 10 | Trogoderma granarium | Beijing Lab rearing | 1.37 | 28 | 24 | 89462 | 77118 |
| 143 | Trogoderma granarium | Beijing Lab rearing | 1.83 | 28 | 25 | 92210 | 80209 |
| 144 | Trogoderma granarium | Beijing Lab rearing | 2.5 | 22 | 18 | 67172 | 58132 |
| 148 | Trogoderma granarium | Beijing Lab rearing | 6.79 | 32 | 29 | 106696 | 88219 |
| 196 | Trogoderma granarium | Beijing Lab rearing | 166.73 | 28 | 24 | 90862 | 75328 |
| 197 | Trogoderma granarium | Beijing Lab rearing | 159.7 | 24 | 21 | 77036 | 64876 |
| 198 | Trogoderma granarium | Beijing Lab rearing | 145.71 | 30 | 26 | 96214 | 79633 |
| 169 | Bactrocera cucurbitae | Beijing Lab rearing | 26.7 | 19.6 | 18 | 65234 | 57356 |
| 170 | Bactrocera cucurbitae | Beijing Lab rearing | 3.89 | 26 | 23 | 86292 | 74433 |
| 171 | Bactrocera cucurbitae | Beijing Lab rearing | 11.53 | 0.13 | 0.057 | 424 | 49 |
| 172 | Bactrocera cucurbitae | Beijing Lab rearing | 8.9 | 32 | 27 | 100670 | 85782 |
| 173 | Bactrocera cucurbitae | Beijing Lab rearing | 33.23 | 34 | 30 | 111070 | 91424 |
| 174 | Bactrocera cucurbitae | Beijing Lab rearing | 4.9 | 28 | 25 | 92402 | 82506 |
| 175 | Bactrocera cucurbitae | Beijing Lab rearing | 13.59 | 30 | 26 | 96954 | 83404 |
| 176 | Bactrocera cucurbitae | Beijing Lab rearing | 35.18 | 22 | 20 | 73186 | 64243 |
| 177 | Bactrocera cucurbitae | Beijing Lab rearing | 59.56 | 30 | 27 | 98028 | 85305 |
| 178 | Bactrocera cucurbitae | Beijing Lab rearing | 5.57 | 30 | 26 | 96282 | 83086 |
| 179 | Bactrocera cucurbitae | Beijing Lab rearing | 9.68 | 24 | 22 | 80230 | 70418 |
| 180 | Bactrocera cucurbitae | Beijing Lab rearing | 14.29 | 28 | 25 | 90368 | 78074 |
| 124 | Solenopsis invicta | Jinghong of Yunnan Province | 0.97 | 24 | 21 | 76194 | 63630 |
| 125 | Solenopsis invicta | Jinghong of Yunnan Province | 1.5 | 40 | 35 | 129366 | 112125 |
| 107 | Dendroctonus pseudotsugae | Taicang of Jiangsu Province | 13.35 | 24 | 20 | 75246 | 63105 |
| 108 | Dendroctonus pseudotsugae | Taicang of Jiangsu Province | 19.44 | 32 | 28 | 104990 | 83408 |
| 109 | Dendroctonus pseudotsugae | Taicang of Jiangsu Province | 9.8 | 30 | 27 | 100354 | 88254 |
| 110 | Dendroctonus pseudotsugae | Taicang of Jiangsu Province | 8.86 | 32 | 28 | 102640 | 85043 |
| 111 | Dendroctonus pseudotsugae | Taicang of Jiangsu Province | 14.49 | 26 | 23 | 85200 | 75105 |
| 72 | Bactrocera dorsalis | Beijing Lab rearing | 5.3 | 30 | 26 | 96956 | 82036 |
| 73 | Bactrocera dorsalis | Beijing Lab rearing | 5.19 | 26 | 23 | 85204 | 72917 |
| 74 | Bactrocera dorsalis | Beijing Lab rearing | 5.85 | 26 | 23 | 86316 | 74316 |
| 75 | Bactrocera dorsalis | Beijing Lab rearing | 3.38 | 26 | 22 | 83080 | 72095 |
| 76 | Bactrocera dorsalis | Beijing Lab rearing | 3.07 | 18.8 | 17 | 62692 | 55309 |
| 1 | Leptinotarsa decemlineata | Urumqi of the Xinjiang Uygur Autonomous Region | 670.47 | 38 | 33 | 123296 | 102809 |
| 2 | Leptinotarsa decemlineata | Urumqi of the Xinjiang Uygur Autonomous Region | 892.25 | 32 | 29 | 107302 | 89862 |
| 3 | Leptinotarsa decemlineata | Urumqi of the Xinjiang Uygur Autonomous Region | 35.93 | 34 | 30 | 112934 | 95863 |
| 4 | Leptinotarsa decemlineata | Urumqi of the Xinjiang Uygur Autonomous Region | 88.16 | 38 | 33 | 124422 | 100516 |
| 5 | Leptinotarsa decemlineata | Urumqi of the Xinjiang Uygur Autonomous Region | 9.69 | 30 | 27 | 98342 | 85244 |
| 6 | Leptinotarsa decemlineata | Urumqi of the Xinjiang Uygur Autonomous Region | 27.69 | 6.4 | 5.7 | 21376 | 18252 |
| 7 | Leptinotarsa decemlineata | Urumqi of the Xinjiang Uygur Autonomous Region | 48.74 | 26 | 23 | 86750 | 73021 |
| 8 | Leptinotarsa decemlineata | Urumqi of the Xinjiang Uygur Autonomous Region | 24.42 | 24 | 22 | 79692 | 69298 |
| 9 | Leptinotarsa decemlineata | Urumqi of the Xinjiang Uygur Autonomous Region | 76.78 | 28 | 25 | 92714 | 80016 |
| 16 | Leptinotarsa decemlineata | Suifenhe of Heilongjiang Province | 23.06 | 38 | 33 | 122058 | 102119 |
| 17 | Leptinotarsa decemlineata | Suifenhe of Heilongjiang Province | 24.67 | 30 | 25 | 96470 | 76947 |
| 18 | Leptinotarsa decemlineata | Suifenhe of Heilongjiang Province | 74.92 | 30 | 27 | 100096 | 82337 |
| 19 | Leptinotarsa decemlineata | Suifenhe of Heilongjiang Province | 74.31 | 36 | 32 | 119348 | 99630 |
| 20 | Leptinotarsa decemlineata | Mishan of Heilongjiang Province | 122.02 | 34 | 29 | 109818 | 92580 |
| 21 | Leptinotarsa decemlineata | Mishan of Heilongjiang Province | 365.71 | 24 | 22 | 79868 | 68195 |
| 22 | Leptinotarsa decemlineata | Mishan of Heilongjiang Province | 256.77 | 26 | 22 | 80878 | 69909 |
| 147 | Leptinotarsa decemlineata | Mishan of Heilongjiang Province | 43.99 | 28 | 25 | 92594 | 77303 |
| 202 | Leptinotarsa decemlineata | Mishan of Heilongjiang Province | 245.92 | 26 | 23 | 83882 | 72339 |
| 203 | Leptinotarsa decemlineata | Mishan of Heilongjiang Province | 203.97 | 28 | 25 | 93658 | 80481 |
| 27 | Henosepilachna vigintioctopunctata | Dongning of Heilongjiang Province | 80.23 | 28 | 25 | 93612 | 79631 |
| 28 | Henosepilachna vigintioctopunctata | Dongning of Heilongjiang Province | 117.05 | 42 | 38 | 140648 | 120691 |
| 29 | Henosepilachna vigintioctopunctata | Dongning of Heilongjiang Province | 69.52 | 34 | 30 | 112460 | 95827 |
| 30 | Henosepilachna vigintioctopunctata | Dongning of Heilongjiang Province | 35.95 | 18.6 | 17 | 61872 | 51118 |
| 31 | Henosepilachna vigintioctopunctata | Hulin of Heilongjiang Province | 93.87 | 28 | 25 | 90406 | 79031 |
| 32 | Henosepilachna vigintioctopunctata | Hulin of Heilongjiang Province | 88.95 | 32 | 29 | 105464 | 89573 |
| 33 | Henosepilachna vigintioctopunctata | Hulin of Heilongjiang Province | 128.1 | 28 | 25 | 91454 | 78455 |
| 34 | Henosepilachna vigintioctopunctata | Hulin of Heilongjiang Province | 126.27 | 24 | 22 | 79250 | 69404 |
| 35 | Henosepilachna vigintioctopunctata | Hulin of Heilongjiang Province | 156.36 | 28 | 25 | 90936 | 79575 |
| 58 | Henosepilachna vigintioctopunctata | Hulin of Heilongjiang Province | 211.32 | 32 | 28 | 105228 | 86584 |
| 59 | Henosepilachna vigintioctopunctata | Hulin of Heilongjiang Province | 572.32 | 28 | 25 | 93512 | 78906 |
| 60 | Henosepilachna vigintioctopunctata | Hulin of Heilongjiang Province | 59.22 | 15.6 | 14 | 52126 | 44679 |
| 61 | Henosepilachna vigintioctopunctata | Hulin of Heilongjiang Province | 36.9 | 26 | 23 | 83838 | 70926 |
| 62 | Henosepilachna vigintioctopunctata | Hulin of Heilongjiang Province | 223.75 | 32 | 28 | 103718 | 87873 |
| 113 | Henosepilachna vigintioctopunctata | Suifenhe of Heilongjiang Province | 31.26 | 34 | 29 | 109096 | 90783 |
| 114 | Henosepilachna vigintioctopunctata | Suifenhe of Heilongjiang Province | 12.2 | 24 | 21 | 77896 | 65920 |
| 115 | Henosepilachna vigintioctopunctata | Suifenhe of Heilongjiang Province | 35.91 | 54 | 47 | 175474 | 147458 |
| 116 | Henosepilachna vigintioctopunctata | Suifenhe of Heilongjiang Province | 29.17 | 19.4 | 18 | 64688 | 55777 |
| 117 | Henosepilachna vigintioctopunctata | Suifenhe of Heilongjiang Province | 7.48 | 30 | 26 | 95840 | 83284 |
| 99 | Bactrocera tau | Beijing Lab rearing | 19.67 | 17.6 | 16 | 58458 | 49831 |
| 100 | Bactrocera tau | Beijing Lab rearing | 13.28 | 28 | 25 | 90384 | 77388 |
| 101 | Bactrocera tau | Beijing Lab rearing | 8.86 | 22 | 20 | 73464 | 62468 |
| 102 | Bactrocera tau | Beijing Lab rearing | 7.59 | 30 | 26 | 94530 | 81460 |
| 103 | Bactrocera tau | Beijing Lab rearing | 19.48 | 30 | 25 | 93796 | 79437 |
| 104 | Bactrocera tau | Beijing Lab rearing | 0.11 | 28 | 25 | 91064 | 77110 |
| 105 | Bactrocera tau | Beijing Lab rearing | 9.87 | 32 | 28 | 103358 | 85488 |
| 106 | Bactrocera tau | Beijing Lab rearing | 69.18 | 32 | 29 | 107058 | 92016 |
| 165 | Bactrocera tau | Beijing Lab rearing | 67.04 | 30 | 26 | 97446 | 82548 |
| 166 | Bactrocera tau | Beijing Lab rearing | 19.55 | 32 | 28 | 101180 | 87621 |
| 167 | Bactrocera tau | Beijing Lab rearing | 11.17 | 26 | 22 | 81050 | 68684 |
| 168 | Bactrocera tau | Beijing Lab rearing | 43.24 | 24 | 21 | 74934 | 64568 |
| 187 | Platypus parallelus | Yangzhou intercepted from the Solomon islands | 1.29 | 36 | 32 | 118686 | 100122 |
| 188 | Platypus parallelus | Yangzhou intercepted from the Solomon islands | 1.77 | 22 | 20 | 73014 | 63471 |
| 189 | Platypus parallelus | Yangzhou intercepted from the Solomon islands | 2.3 | 30 | 27 | 98414 | 83773 |
| 118 | Cydia pomonella | Dongning of Heilongjiang Province | 11.55 | 26 | 23 | 84282 | 71440 |
| 119 | Cydia pomonella | Dongning of Heilongjiang Province | 4.18 | 28 | 25 | 91814 | 78652 |
| 120 | Cydia pomonella | Dongning of Heilongjiang Province | 8.72 | 26 | 22 | 83478 | 69046 |
| 121 | Cydia pomonella | Dongning of Heilongjiang Province | 5.46 | 32 | 27 | 101484 | 81648 |
| 11 | Cydia pomonella | Mudanjiang of Heilongjiang Province | 3.9 | 24 | 22 | 79338 | 69044 |
| 12 | Cydia pomonella | Mudanjiang of Heilongjiang Province | 13.22 | 24 | 20 | 74196 | 64095 |
| 13 | Cydia pomonella | Mudanjiang of Heilongjiang Province | 2.75 | 28 | 24 | 89158 | 78374 |
| 14 | Cydia pomonella | Mudanjiang of Heilongjiang Province | 16.34 | 32 | 28 | 103688 | 87260 |
| 15 | Cydia pomonella | Mudanjiang of Heilongjiang Province | 7.21 | 34 | 30 | 110938 | 82637 |
| 63 | Cydia pomonella | Urumqi of the Xinjiang Uygur Autonomous Region | 15.84 | 28 | 24 | 89646 | 76066 |
| 64 | Cydia pomonella | Urumqi of the Xinjiang Uygur Autonomous Region | 24.18 | 28 | 24 | 90044 | 77669 |
| 65 | Cydia pomonella | Urumqi of the Xinjiang Uygur Autonomous Region | 4.85 | 18.2 | 17 | 60834 | 51906 |
| 201 | Cydia pomonella | Urumqi of the Xinjiang Uygur Autonomous Region | 12.34 | 30 | 26 | 97020 | 80077 |
| 82 | Cydia pomonella | Urumqi intercepted from Kazakhstan | 2.62 | 32 | 27 | 101214 | 80334 |
| 83 | Cydia pomonella | Urumqi intercepted from Kazakhstan | 2.17 | 36 | 32 | 116344 | 101987 |
| 84 | Cydia pomonella | Urumqi intercepted from Kazakhstan | 5.49 | 32 | 28 | 104646 | 85988 |
| 85 | Cydia pomonella | Urumqi intercepted from Kazakhstan | 1.07 | 36 | 31 | 115958 | 94165 |
| 86 | Cydia pomonella | Urumqi intercepted from Kazakhstan | 3.14 | 32 | 28 | 105382 | 86733 |
| 129 | Cydia pomonella | Korla of the Xinjiang Uygur Autonomous Region | 5.31 | 22 | 20 | 73484 | 63733 |
| 130 | Cydia pomonella | Korla of the Xinjiang Uygur Autonomous Region | 6.33 | 30 | 26 | 94500 | 80533 |
| 131 | Cydia pomonella | Korla of the Xinjiang Uygur Autonomous Region | 5.19 | 30 | 26 | 97716 | 82687 |
| 132 | Cydia pomonella | Korla of the Xinjiang Uygur Autonomous Region | 2.84 | 20 | 18 | 66398 | 56300 |
| 133 | Cydia pomonella | Korla of the Xinjiang Uygur Autonomous Region | 5.3 | 28 | 25 | 91478 | 77508 |
| 66 | Cydia pomonella | Ili of the Xinjiang Uygur Autonomous Region | 6.86 | 42 | 37 | 137196 | 117395 |
| 126 | Eriosoma lanigerum | Mudanjiang of Heilongjiang | 3.76 | 32 | 27 | 102296 | 84011 |
| 127 | Eriosoma lanigerum | Mudanjiang of Heilongjiang | 4.3 | 30 | 26 | 95052 | 79658 |
| 128 | Eriosoma lanigerum | Mudanjiang of Heilongjiang | 4.21 | 58 | 50 | 188188 | 156747 |
| 145 | Eriosoma lanigerum | Mudanjiang of Heilongjiang | 1.88 | 24 | 22 | 78618 | 68570 |
| 157 | Eriosoma lanigerum | Mudanjiang of Heilongjiang | 0.97 | 18.4 | 17 | 61242 | 53150 |
| 158 | Eriosoma lanigerum | Mudanjiang of Heilongjiang | 0.42 | 26 | 23 | 83436 | 71607 |
| 159 | Eriosoma lanigerum | Mudanjiang of Heilongjiang | 0.68 | 22 | 19 | 70770 | 59724 |
| 160 | Eriosoma lanigerum | Mudanjiang of Heilongjiang | 0.74 | 30 | 27 | 100088 | 83948 |
| 136 | Lymantria dispar | Beijing Lab rearing | 64.38 | 28 | 25 | 89762 | 78769 |
| 137 | Lymantria dispar | Beijing Lab rearing | 50.97 | 16 | 15 | 53134 | 46642 |
| 184 | Dysmicoccus neobrevipes | Beijing Lab rearing | 41.5 | 30 | 26 | 96352 | 84724 |
| 185 | Dysmicoccus neobrevipes | Beijing Lab rearing | 54.42 | 36 | 32 | 119364 | 99674 |
| 186 | Dysmicoccus neobrevipes | Beijing Lab rearing | 70.17 | 30 | 26 | 94100 | 83187 |
| 25 | Brontispa longissima | Haikou of Hainan Province | 6.32 | 36 | 31 | 115420 | 98163 |
| 26 | Brontispa longissima | Haikou of Hainan Province | 8.1 | 28 | 24 | 87624 | 76870 |
| 57 | Brontispa longissima | Haikou of Hainan Province | 6.32 | 34 | 30 | 111624 | 90538 |
| 23 | Opisina arenosella | Haikou of Hainan Province | 53.17 | 30 | 26 | 96272 | 82046 |
| 24 | Opisina arenosella | Haikou of Hainan Province | 18.85 | 28 | 25 | 93214 | 80231 |
| 56 | Opisina arenosella | Haikou of Hainan Province | 17.81 | 28 | 25 | 91630 | 80694 |
| 134 | Sitophilus zeamais | Shanghai | 5.01 | 32 | 28 | 105074 | 89530 |
| 135 | Sitophilus zeamais | Shanghai | 5.72 | 22 | 18 | 67132 | 58226 |
| 204 | Sitophilus zeamais | Shanghai | 12.53 | 30 | 26 | 96096 | 80358 |
| 67 | Ips typographus | Taicang of Jiangsu Province | 3.97 | 36 | 32 | 118542 | 97859 |
| 68 | Ips typographus | Taicang of Jiangsu Province | 1.74 | 32 | 27 | 101756 | 82250 |
| 69 | Ips typographus | Taicang of Jiangsu Province | 0.83 | 32 | 29 | 105272 | 92676 |
| 70 | Ips typographus | Taicang of Jiangsu Province | 1.56 | 24 | 20 | 74194 | 63485 |
| 71 | Ips typographus | Taicang of Jiangsu Province | 1.17 | 32 | 28 | 103474 | 87222 |
| 77 | Ips typographus | Huangdao port intercepted from Czech | 3.25 | 26 | 22 | 80624 | 69541 |
| 78 | Ips typographus | Huangdao port intercepted from Czech | 2.62 | 30 | 26 | 94072 | 81640 |
| 79 | Ips typographus | Huangdao port intercepted from Czech | 3.7 | 20 | 18 | 66570 | 55117 |
| 80 | Ips typographus | Huangdao port intercepted from Czech | 2.97 | 38 | 34 | 127444 | 108081 |
| 81 | Ips typographus | Huangdao port intercepted from Czech | 2.35 | 30 | 27 | 99202 | 80573 |
| 138 | Ips typographus | Suifenhe of Heilongjiang Province | 2.85 | 26 | 23 | 84326 | 73084 |
| 141 | Ips typographus | Suifenhe of Heilongjiang Province | 3.15 | 24 | 20 | 74420 | 65587 |
| 142 | Ips typographus | Suifenhe of Heilongjiang Province | 2.14 | 24 | 21 | 74728 | 65440 |
| 149 | Ips typographus | Suifenhe of Heilongjiang Province | 3.02 | 26 | 24 | 86452 | 76110 |
| 150 | Ips typographus | Suifenhe of Heilongjiang Province | 3.96 | 34 | 29 | 108618 | 91677 |
| 151 | Ips typographus | Suifenhe of Heilongjiang Province | 29.42 | 24 | 20 | 73720 | 63867 |
| 152 | Ips typographus | Suifenhe of Heilongjiang Province | 6.52 | 26 | 22 | 82066 | 70556 |
| 181 | Carpomya vesuviana | Ili of the Xinjiang Uygur Autonomous Region | 6.61 | 32 | 28 | 103152 | 88382 |
| 182 | Carpomya vesuviana | Ili of the Xinjiang Uygur Autonomous Region | 0.56 | 32 | 29 | 105620 | 91904 |
| 183 | Carpomya vesuviana | Ili of the Xinjiang Uygur Autonomous Region | 2.52 | 24 | 21 | 77936 | 68116 |
