## Supplementary figures and images for "First Glimpse of Gut Microbiota of Quarantine Insects in China"

### Supplemental Figure 1

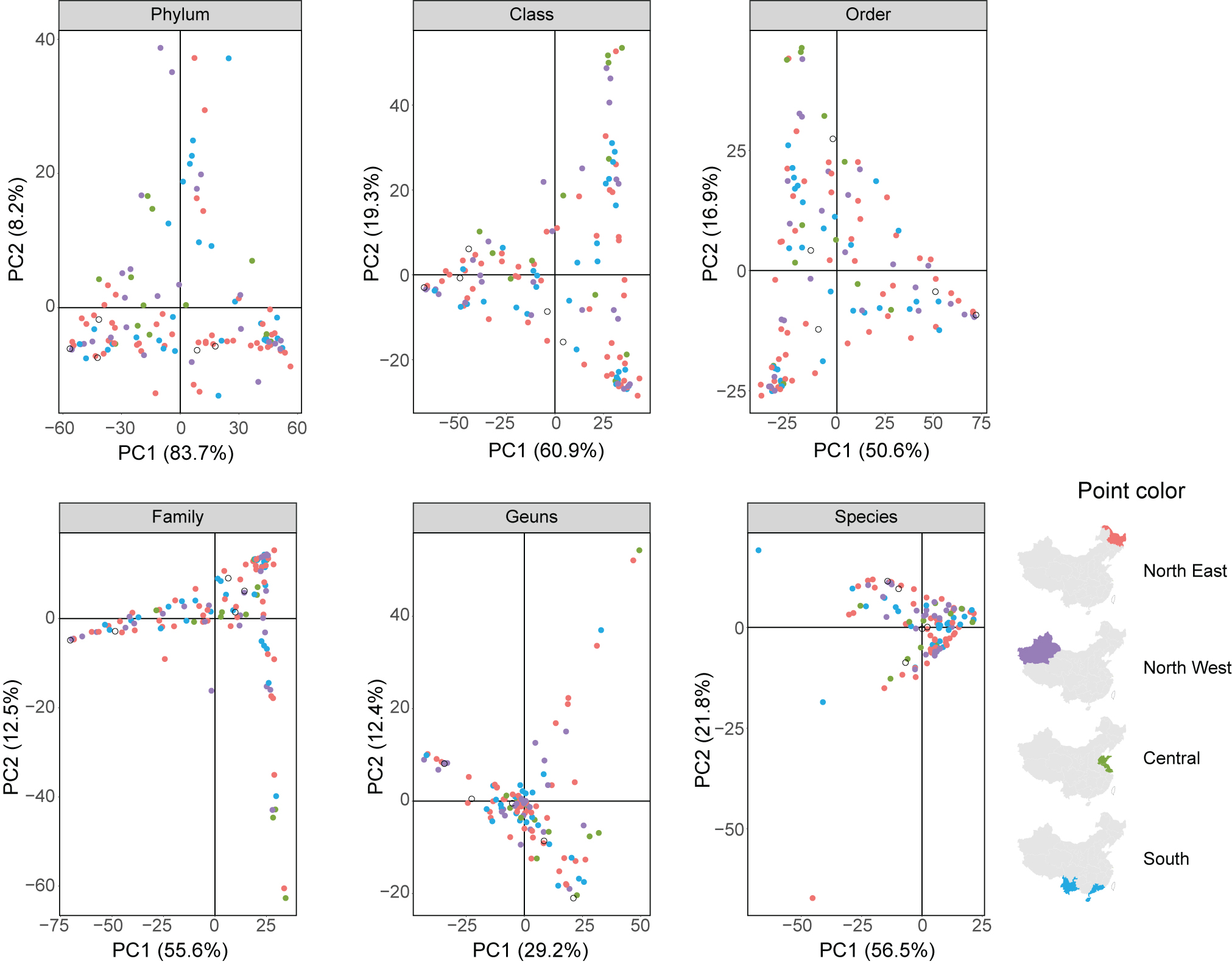

### Supplemental Figure 2

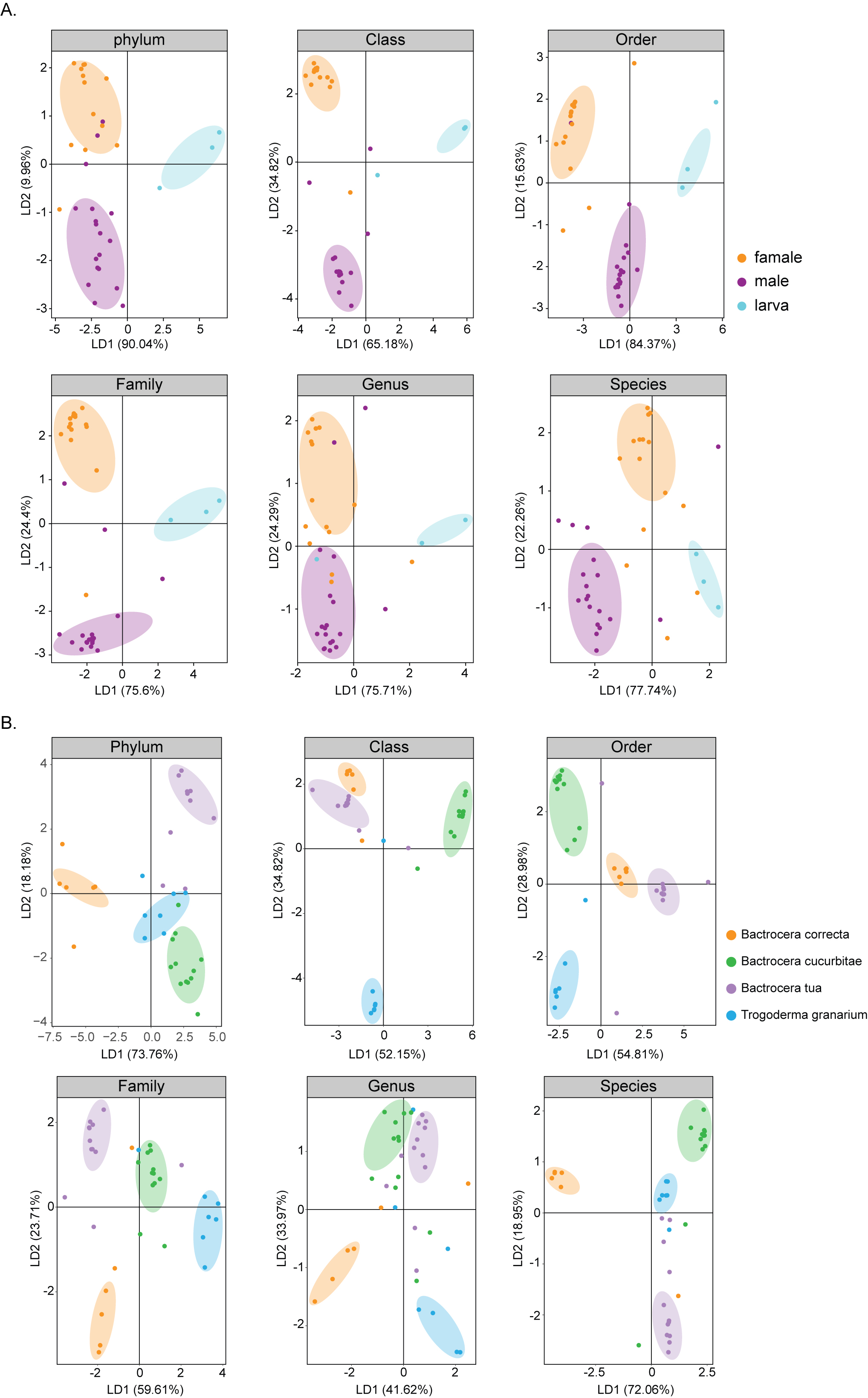

### Supplemental Figure 3

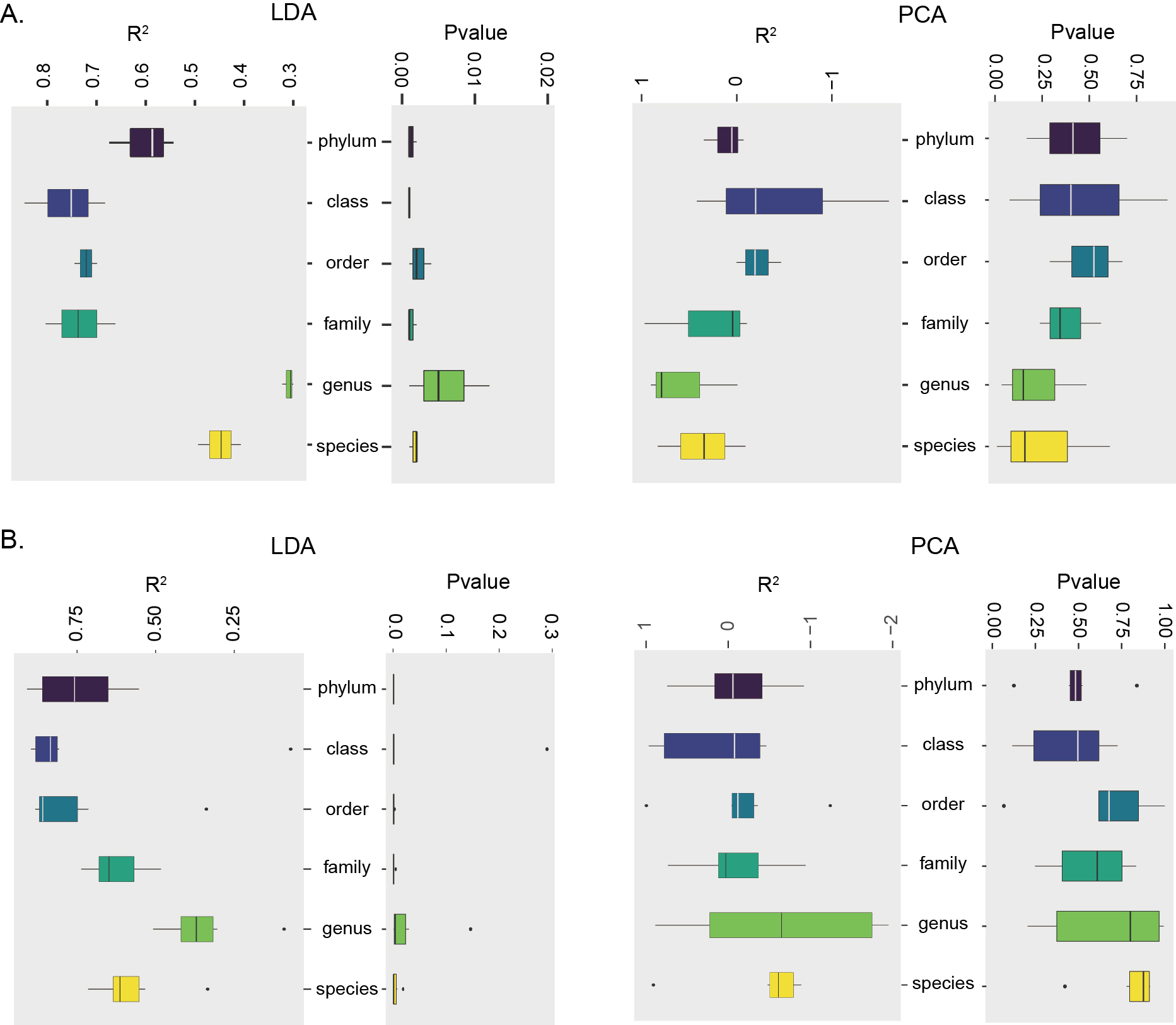

### Supplemental Figure 4

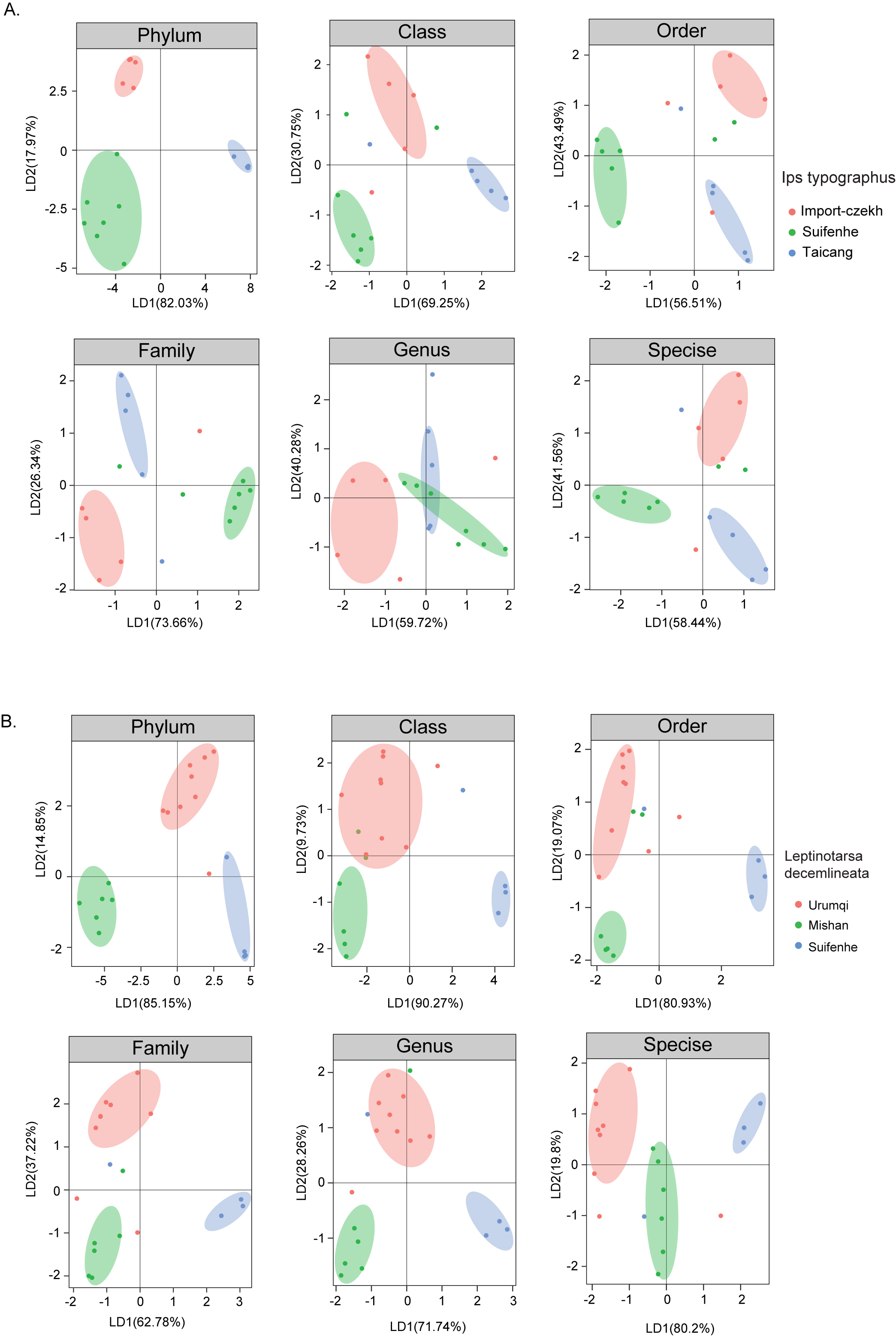

### Supplemental Figure 5

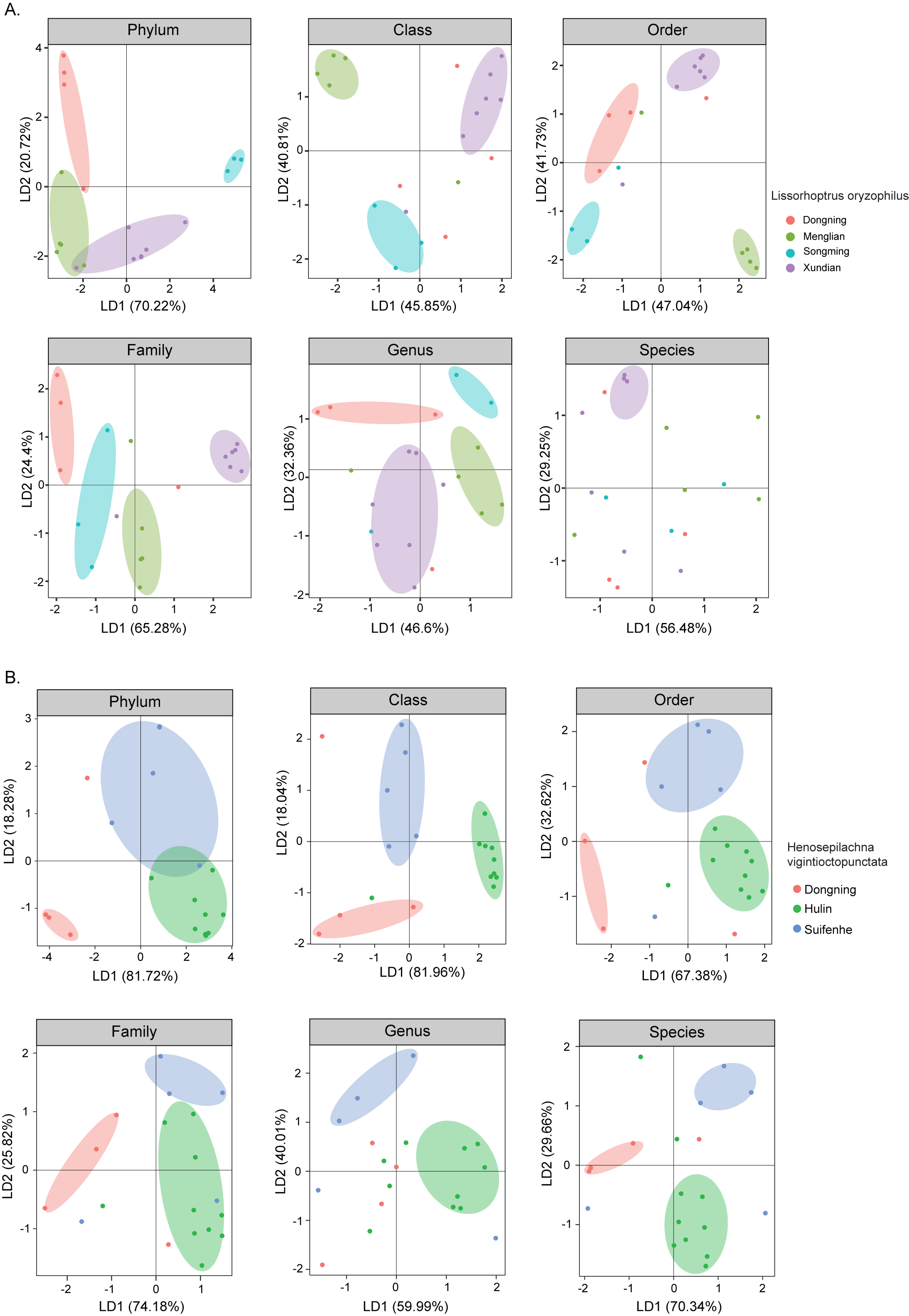

### Supplemental Figure 6

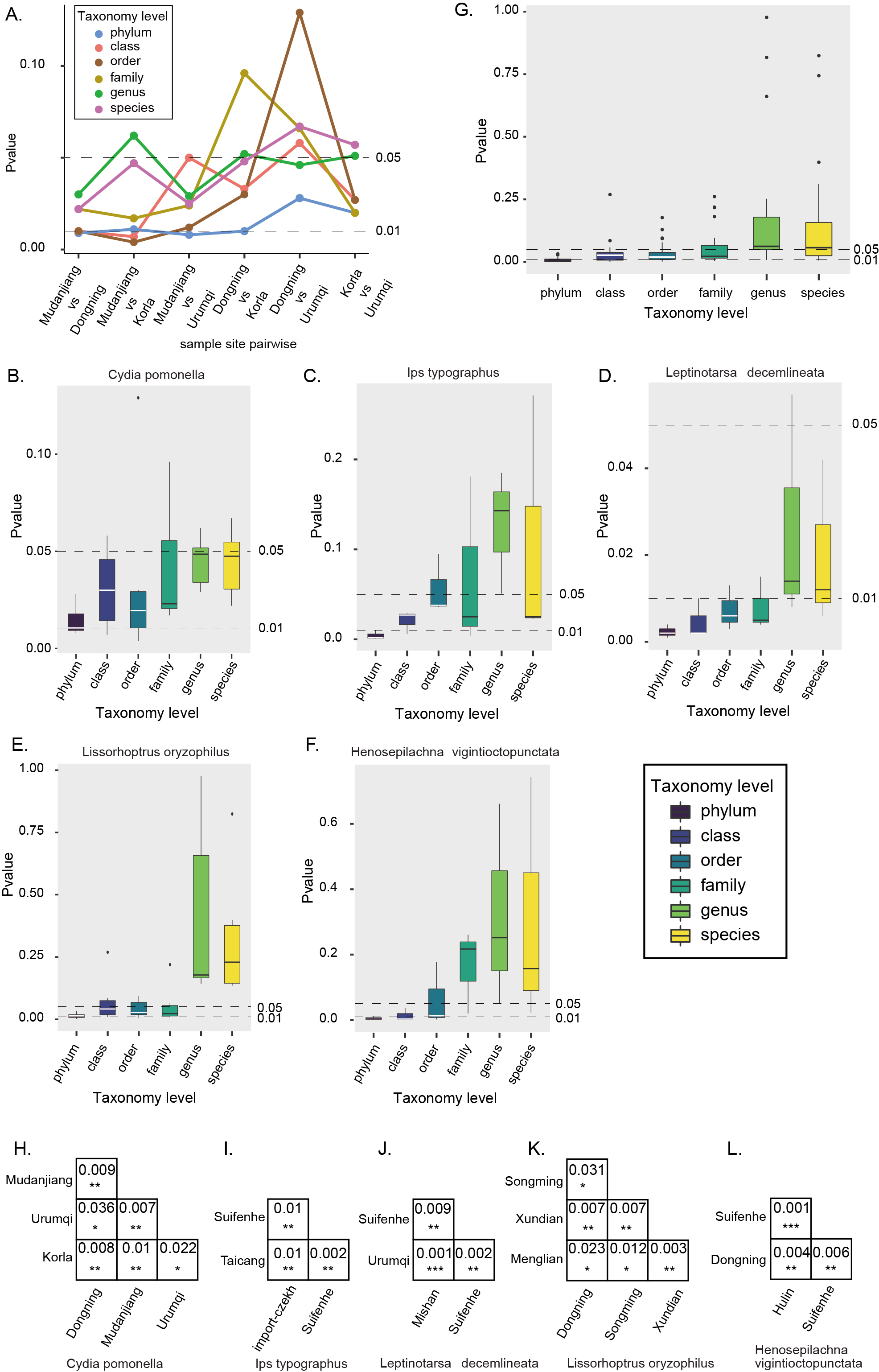
